# VIDEO - Visual Integration of Drosophila Enhancer Organization: A tool for integrating and visualizing chromatin accessibility, in vivo transcription factor binding and motif occurrence in tissue-specific differentially expressed genes

**DOI:** 10.1101/2025.10.27.684897

**Authors:** Vidya Ajay, Nathaniel Laughner, Patrick Cahan, Deborah J. Andrew

**Affiliations:** Department of Cell Biology, 725 N. Wolfe St., Hunterian G1, Johns Hopkins University School of Medicine, Baltimore, MD 21205, USA; Department of Biomedical Engineering, 725 N. Wolfe St., Hunterian G1, Johns Hopkins University School of Medicine, Baltimore, MD 21205, USA

**Keywords:** Assay for Transposase-Accessible Chromatin with high-throughput sequencing (ATAC seq), Chromatin Immunoprecipitation Sequencing (ChIP-Seq), DNA binding motifs, promoter proximal enhancers, single cell RNA-Seq (scRNA-Seq)

## Abstract

Dissecting gene regulation today relies on many genomic assays – including transcriptional output from RNA-seq, chromatin accessibility from ATAC-seq, and transcription factor (TF) binding from ChIP-seq. Whereas numerous tools exist for each modality and some integrate data across modalities, few allow researchers to interactively explore and visualize how TF binding motifs intersect with transcriptional activity and chromatin accessibility in a tissue-specific context. Here we introduce VIDEO (<u>V</u>isual <u>I</u>ntegration of <u>D</u>rosophila <u>E</u>nhancer <u>O</u>rganization), a web-based analysis tool that enables visualization of conserved TF binding motifs within proximal promoters of genes differentially expressed in specific tissues. Starting with gene lists derived from in situ hybridization, microarray, and/or scRNA-seq studies of WT or mutant samples, one can identify the TFs expressed in each tissue and learn if and where the consensus binding motifs for those TFs are found within the proximal enhancers of a custom gene set. This pipeline also allows for coincident visualization of active chromatin, as determined from ATAC-seq data, and for the visualization of DNA binding data from ChIP-seq datasets for specific TFs. To demonstrate its utility, we apply VIDEO to the well-characterized regulatory system of CrebA and the secretory pathway in the *Drosophila* salivary gland. We also explore a lesser-known system in the embryonic hindgut to show how utilization of this tool can serve to generate hypotheses regarding regulatory interactions.

## Introduction

The application of recent advances in high-throughput sequencing have revealed large sets of differentially expressed (DE) genes from multiple *Drosophila* cell types at distinct developmental stages and under a variety of mutant conditions (Karaiskos *et al*. 2017; Calderon *et al*. 2022; Seroka *et al*. 2022; Sakaguchi *et al*. 2023; Peng *et al*. 2024; Jackson *et al*. 2025; Wang *et al*. 2025). Related technologies have likewise generated extensive *Drosophila* chromatin accessibility and transcription factor (TF) binding datasets with associated binding site consensus motif data (Bergman *et al*. 2005; Zhu *et al*. 2011; Shazman *et al*. 2014; Nitta *et al*. 2015; Kudron *et al*. 2024; Rauluseviciute *et al*. 2024). These motifs, 6-12 bp long sequences recognized by specific transcription factors, are a critical component of gene regulatory networks, and mapping their patterning across the genome is crucial to formulating hypotheses about TF-target gene interactions. A challenge that remains is connecting these datasets to identify which TFs may regulate a given set of DE genes and where their binding motifs occur in open chromatin.

Existing workflows for motif analysis extract peak regions from ATAC-seq (open chromatin) and ChIP-seq (TF occupancy) data, convert them to FASTA sequences, and run motif discovery tools such as MEME (Bailey and Elkan 1994), HOMER (Heinz *et al*. 2010) and RSAT (Thomas-Chollier *et al*. 2012). Single-cell RNA-seq further expands possible motif-analysis inputs by generating lists of genes of interest from differential expression analysis. For all inputs, users must retrieve the peak or promoter/enhancer sequences, wrangle multiple file formats, and merge motif hits with chromatin or binding tracks using command-line tools (e.g. bedtools) and write custom scripts for visualization. This fragmented, multi-step workflow and the lack of user-friendly visualization tools often leaves motif data under-utilized in the context of gene regulatory network studies, and there is still a lack of an intuitive way to visualize and explore TF-motif landscapes in their native genomic context.

Existing visualization tools address parts of the pipeline; MEME-Suite (Bailey *et al*. 2015) allows for a basic position display for motifs it discovers but does not accommodate externally defined motifs. TFMotifView (Leporcq *et al*. 2020) allows browsing of a pre-computed motif library in user-defined genomic regions, but it does not take as input user-generated motifs in standard formats (PFM, PWM, PCM, IUPAC as described in **Supplemental Table S1**) and it does not support the latest *Drosophila* genome assemblies. Additionally, the output of these existing tools fails to utilize the potential of overlaying TF motifs with user-discovered gene lists and sequencing data.

To address these shortcomings, we have built VIDEO, a web-based platform designed specifically for use with *Drosophila*, with a streamlined, user-friendly graphical interface that requires no programming knowledge. VIDEO integrates processed ATAC-seq and ChIP-seq peak information into signal tracks with genomic input, integrates motif discovery while also accepting user-defined motifs in any common format, and displays motif hits alongside genomic signal overlays. VIDEO allows users to rapidly explore many transcriptional regulation hypotheses and questions such as: Are the motifs of the TF of interest enriched in the user-provided list? Do they occur more densely in the list compared to a control group? Which isoform/transcription start site (TSS) is the richest in the TF motif? Do motif hits occur in regions of accessible chromatin? Which of these hits are supported by DNA binding data? Currently, there is no single public resource that can be mined to address all these questions. VIDEO also serves as a data bank for users who want to extract processed scRNA sequencing data for *Drosophila* embryos (stage 10-16, (Peng *et al*. 2024) and find TF motifs of interest, enabling both an end-to-end pipeline for initial hypothesis generation, and a much more detailed and focused pipeline for exploring motif occurrences in user-generated gene lists with supporting ChIP-seq and ATAC-seq data. VIDEO can be accessed through this link: VIDEO-motif-tool.

## Materials and Methods

### Input data and preprocessing

User inputs consist of either a gene list of FlyBase IDs or official gene symbols or a BED file with genomic peak regions. In the “compare” mode users can input two gene lists with distinct labels and perform the same downstream analysis. For gene-list inputs, transcript annotations for all isoforms and ±500 bp (or a user-defined window size up to 2 kb) flanking sequences around each TSS were indexed from a copy of the *Drosophila melanogaster* reference genome (dm6) FASTA using pyfaidx (v0.8.1.4). For BED-file inputs, peak coordinates were used directly to extract the corresponding FASTA sequences.

### Motif scanning

User-defined motifs (PFM, PWM, PCM, or IUPAC formats) were scanned against the extracted sequences using FIMO (v4.12.0; (Grant *et al*. 2011)) with a default significance threshold of p < 1e-4. A custom Python wrapper launched FIMO as a subprocess, parsed its output, and collated motif hits into a unified table recording genomic coordinates, strand, p-values, q-values, and matched sequences.

### Visualization of motif occurrences

Motif occurrences were plotted with Plotly (v6.2.0) in python. For each gene or peak region, hits were positioned relative to the TSS (zero), spanning ±500 bp by default. The opacity of each hit reflects its FIMO match score, with darker hits indicating a stronger match. Hits with negative log-likelihood ratio scores were excluded, as they are more likely to represent random matches and correspond to higher p-values. Motif hits are ordered either by the user’s input list or by total motif count or mean match score, as selected in the interface. The plot is interactive and all plotting defaults are configurable in the web user interface (UI). Match stringency is an adjustable variable and users can further adjust the p-values for each motif specifically.

### Data Integration

To filter motif hits by chromatin context, ATAC-seq and ChIP-seq peaks were integrated via Pybedtools (v0.12.0) calling bedtools intersect with a minimum overlap of 50% of the motif length.

### BigWig Track Overlay

To visualize chromatin accessibility and transcription factor occupancy in the same genomic window as motif hits, BigWig tracks from ATAC-seq or ChIP-seq experiments were uploaded. For each selected peak or gene, coverage values are extracted from the corresponding chromosome interval using pyBigWig (v0.3.22). Coverage is plotted in Plotly alongside motif positions, with the x-axis centered on the peak midpoint or TSS and consistent y-axis scaling across all uploaded tracks to facilitate comparison. Motif hits from the motif occurrence plot are overlaid in the top panel and strand direction is accounted for.

### VIDEO implementation

VIDEO was implemented using FastAPI (v0.95.2) backend and a React.js (v18.2.0) frontend. Core dependencies include FIMO (v4.12.0), pyfaidx (v0.5.9), Pybedtools (v0.12.0), and Plotly (v6.2.0). The application was deployed on an EC2 Instance with Redis as broker. Reference genome FASTA and GTF annotation (dm6, release 6.45) were downloaded from FlyBase; motif libraries were bulk downloaded from JASPAR (Rauluseviciute *et al*. 2024).

### Data and code availability

All the code used for developing the tool is available at https://github.com/vidyaajay1/VIDEO-motif-tool.

### Embryo immunostaining

Embryos were fixed and immunostained as described (Reuter *et al*. 1990). Rat *α*Retn (Haberman *et al*. 2003) was used at a 1/20,000 dilution. Biotin-conjugated Donkey *α*Rat secondary was used at a dilution of 1:2000.

## Results

VIDEO presents an interactive, browser-based overview designed to guide users from raw inputs to interpretable motif visualizations without the need for multi-tool command-line operations. Here, we provide a flowchart that can be used as a guide for new users (**Fig 1**). Upon starting the application, the landing page has a collapsible sidebar navigator that has two functional pages “Motif Viewer” and “TF Finder” and a tutorial page with instructions (**Fig 2A**). The Motif Viewer has the pipeline for generating the motif occurrence plot, while TF Finder is a helper page with functions intended to provide users with a starting gene list and allow discovery of TFs and motifs of interest.

**Figure 1.**
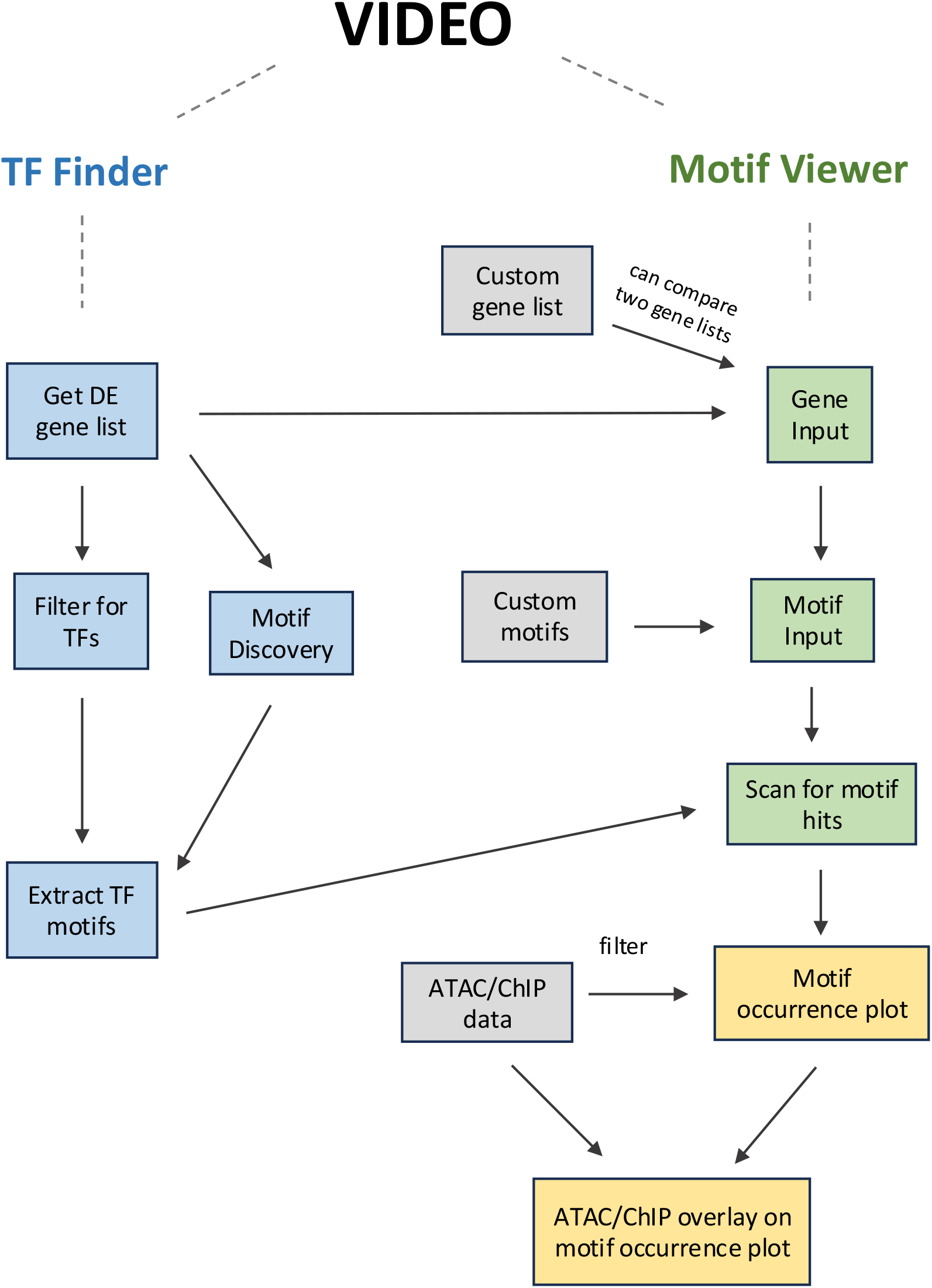
Flowchart of many possible pipelines using VIDEO. TF Finder and Motif Viewer are the two pages in VIDEO. Each blue and green box represents a module in the page In Motif Viewer (green), these modules are presented as navigable cards, and in TF Finder (blue) they are shown one below another in the same page. Gray boxes represent user-provided data (not native to VIDEO). Yellow boxes represent final outputs from VIDEO. Arrows represent a sequential order of execution of the modules. Arrows start from a module that gives an output and go towards a module that can use this output as its input. Dashed lines represent a hierarchical connection between the pages and their modules.

**Figure 2.**
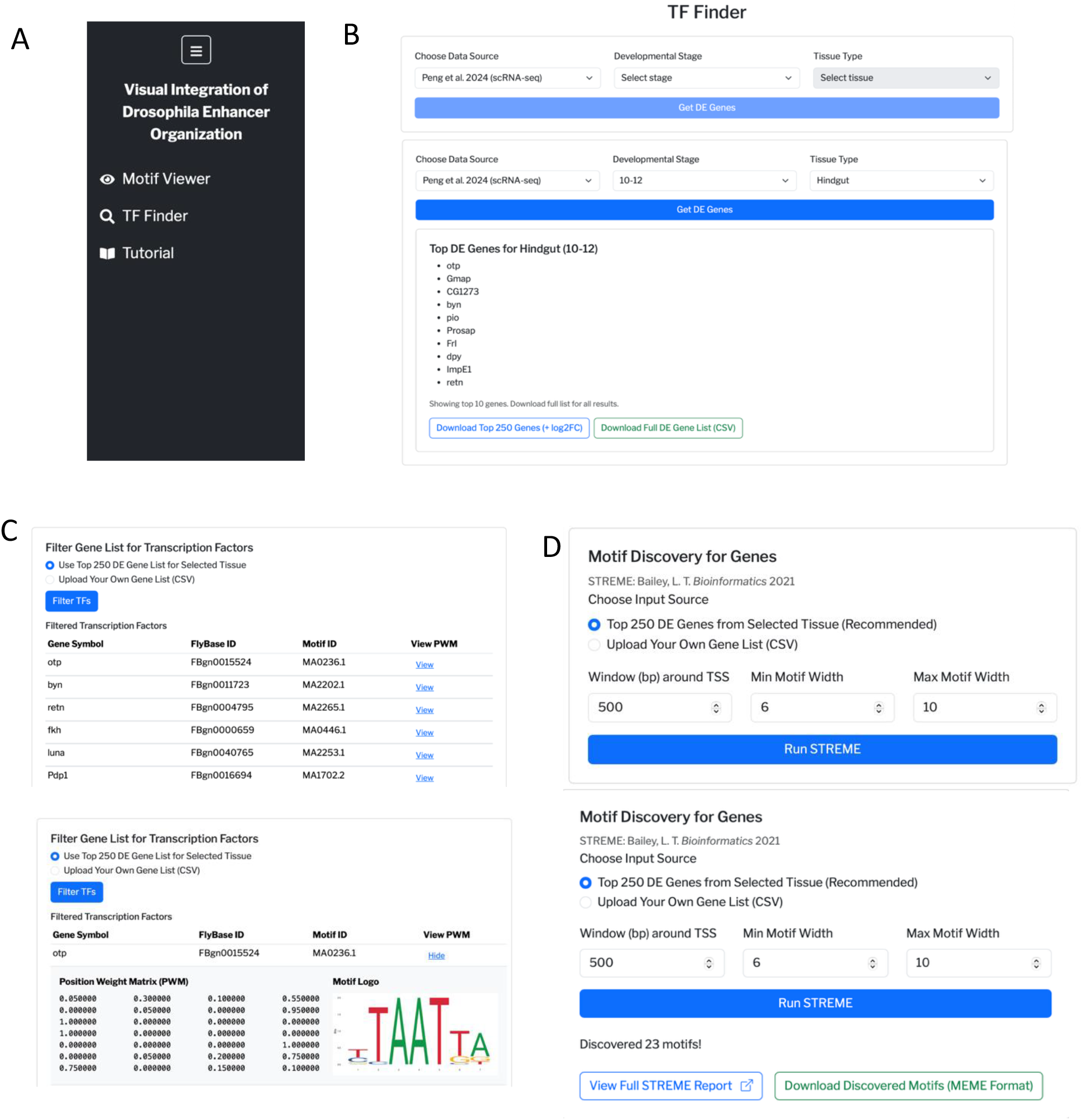
Steps for Using TF finder. **(A)** The collapsible sidebar navigator in VIDEO with two functional pages and a tutorial page. **(B)** Getting DE gene lists from TF Finder. Top: before choosing a developmental stage and tissue type. Bottom: after making the selections and pressing Get DE genes. **(C)** Filtering gene lists for TFs. Top: After pressing Filter TFs, Bottom: Motif details after pressing View PWM **(D)** Discovering motifs from gene lists. Top: Choose parameters and upload gene list before running STREME. Bottom: After pressing Run STREME, with link to outputs and a button to download the discovered motifs.

### 1. Using TF Finder to acquire gene and motif data Identification of gene sets of interest

There are multiple ways a user can acquire a gene list of interest. These include, but are not limited to, mining transcriptomic databases such as BDGP (Tomancak *et al*. 2007) for genes differentially expressed (DE) in a specific tissue, extracting tissue-specific DE genes from scRNA-seq data, and identifying TF-dependent gene sets from comparisons of WT and mutant scRNA-seq, microarray, and/or in situ data. Users are encouraged to rank their gene lists in order of most important to least if there are available metrics such as p-values or log fold change expression, because VIDEO can retain the input order while plotting. To facilitate very straightforward explorations on DE gene sets, TF Finder contains a curated set of DE genes for two stages of development of the *Drosophila* embryo from Peng et. al 2024 single cell RNA sequencing data (**Fig 2B**). Users can choose the stage of development and the tissue, and download a list of the top DE genes in that tissue.

#### Identify all TFs in a gene list

TF Finder has a helper function that filters any gene list for transcription factors (as reported in FlyBase (Ozturk-Colak *et al*. 2024)). Users can identify TFs differentially expressed in their tissue of interest by choosing the “Use Top 250 DE Genes for Selected Tissue” option or upload their own gene list to extract the TFs in it (**Fig 2C**). Once the gene list is filtered for TFs, VIDEO also displays the consensus motifs for each TF (if available) in the JASPAR database with their motif ID, PWM and logo.

#### Identification of TF motifs or DE gene conserved motifs

If the user does not have a TF motif of interest already, or if the TF of interest does not have any reported motifs shown in VIDEO, they can make use of the in-built “Motif Discovery for Genes” function (**Fig 2D**). Again, the user can select the “Top 250 DE Genes for Selected Tissue” option or upload their own gene list. Once the gene list is uploaded, the user can set a promoter region flanking the TSS and the minimum and maximum motif width. If nothing is specified, the program will default to its preset values (6 bp to 10 bp width). On clicking the “Run STREME” button, the motif discovery program will output a link to view and download the motifs that are enriched in the list of genes. Once the motifs of interest are decided, it is recommended that the user keep the motifs in PWM format (MEME format). If the motifs were chosen from VIDEO, the displayed PWM can directly be copied to a clipboard and pasted as motif input.

### 2. Using Motif Viewer to create motif hit plots

#### Setting up genomic input

This is the landing page of VIDEO, and in the regular mode users can upload their gene list or a BED file with genomic regions of interest (**Fig 3A**). In the compare mode, users are required to upload two gene lists and give each list a unique name, as elaborated further in the case study later. Once the gene lists are uploaded, the next step is to choose a promoter window size flanking the TSS of each gene. If a gene has multiple TSSs, all unique TSSs will be considered and annotated with the transcript ID. If no choice is made, the program will default to the preset 500 bp upstream and downstream. The last step is to click on the “Process Genomic Input” button, which validates the gene list and fetches all the transcript annotations and sequences for downstream processing.

**Figure 3.**
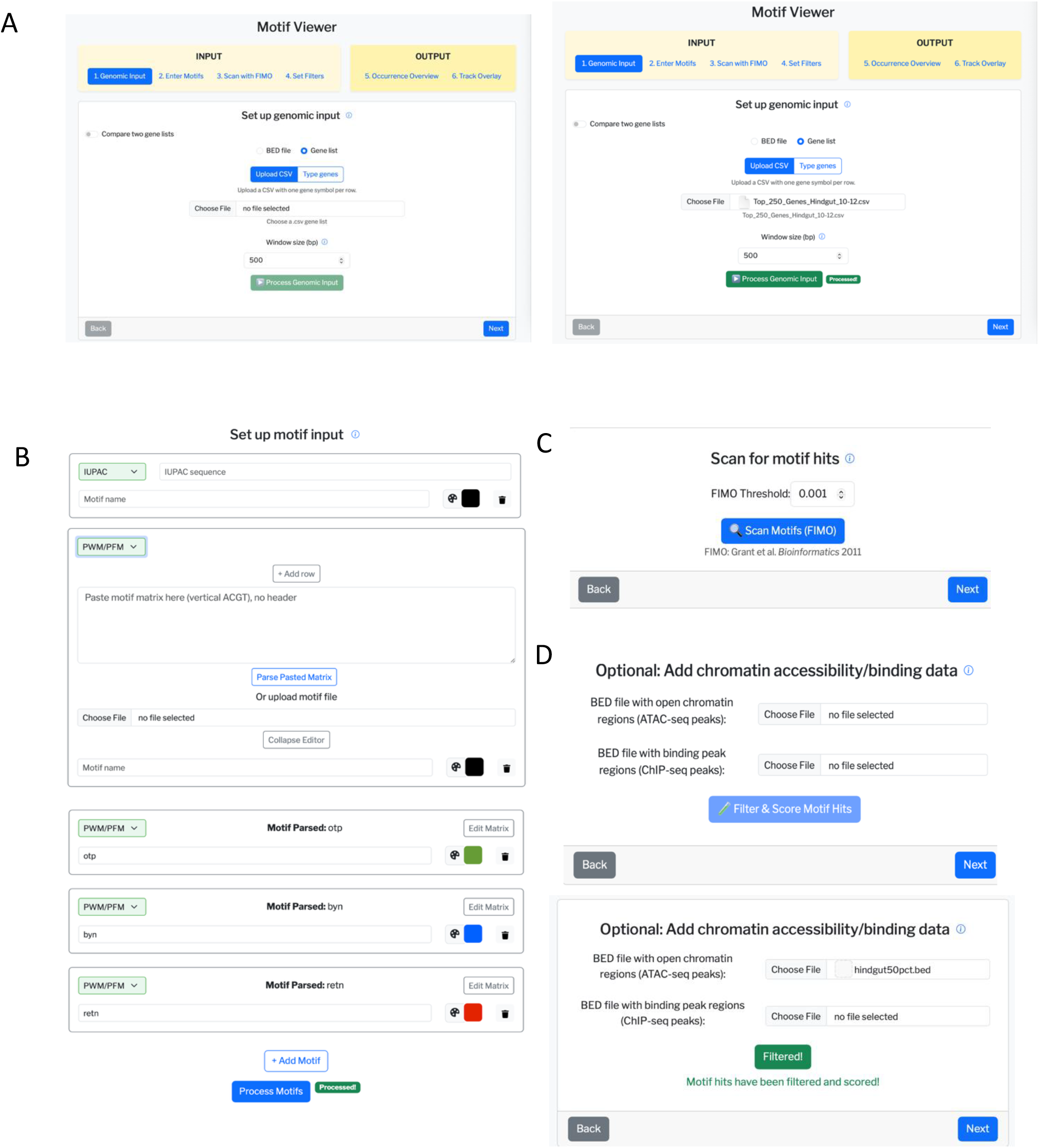
Steps for Using Motif Viewer. **(A)** Set up genomic input. Left: Before uploading gene list, Right: After uploading data, choosing window size and pressing Process Genomic Input. **(B)** Set up motif input. Top: Initial window, Middle: Window after selecting the PWM/PFM format. Paste the motif matrix, press Parse Pasted Matrix and give the motif a name and color. Bottom: After three motifs are added and Process Motifs is pressed. **(C)** Scan for motif hits. User sets the p-value threshold and presses Scan Motifs (FIMO). **(D)** Add ATAC/ChIP-seq data if available. Top: Before uploading data, Bottom: After uploading ATAC data (Wang et al. 2025) and clicking on the Filter and Score Motif Hits button.

#### Setting up motif input

The motif input page requires users to drop in their motifs in any of three formats (IUPAC consensus sequence, PWM, PFM, or PCM), give the motif a name, and choose a color for it for the motif occurrence plot (**Fig 3B**). If users are dropping in PWM, PFM or PCM, they can either upload it as a.txt file or copy and paste the matrix and then click “Parse Pasted Matrix”. Once the matrix is parsed and all the motifs are input, clicking the “Process Motifs” button will both validate the motifs and process it for the plot.

#### Scanning for motif occurrences

After the gene list and motif data are given to the program, it is ready for scanning the gene list for motif occurrences. Users can choose the overall p-value threshold for the motif scanning algorithm (default is 0.001) (**Fig 3C**). Once the p-value is set, the user should click on the “Scan FIMO” button to scan for motif hits.

#### Integrating ATAC-seq and ChIP-seq data

If the user has supporting chromatin accessibility and/or in vivo TF binding data for their gene lists, they can upload the BED files with the respective peaks into VIDEO. This is the fourth window in VIDEO and is completely optional – click on the “Next” button to proceed to the plot if you don’t have this data (**Fig 3D**). VIDEO will use the ATAC and ChIP data as an option to filter the motif occurrences for hits that occur only in “open chromatin” or hits in “ChIP peak” regions and the filters can be turned on and off during plotting. For this demonstration, we mined scATAC data from (Wang *et al*. 2025) for the hindgut tissue in stages 10-12. Once the data are uploaded, the user should press the “Filter Motif Hits” button and then proceed to the next page.

#### Adjusting the plot settings

This window is an array of configurable plot settings and a motif occurrence plot below it (**Fig 4**). If the users chose to upload ATAC/ChIP data they can toggle the filters on and off by checking the respective options under the “Show only:” heading. The motif occurrence plot will default to the order of the input gene list and will show all the transcripts of each gene, but the user has the option to customize this. First, go to the “Sort by motif:” heading and select the motif you want to prioritize. This will display three options, the first two of which are metrics you need to provide for the sorting. Checking “Number of Hits” will sort the gene list by the number of hits for the selected motif, and checking “Motif Match Score” will sort the gene list by the motif match score of the best scoring motif hit for each gene. By default, all the transcripts of a gene will be clustered together with the top scoring transcript, and the user has the option to choose “Show only best transcripts for selected motif hit”, which will show only the top ranking TSS for each gene based on the metrics the user selected for ranking.

**Figure 4.**
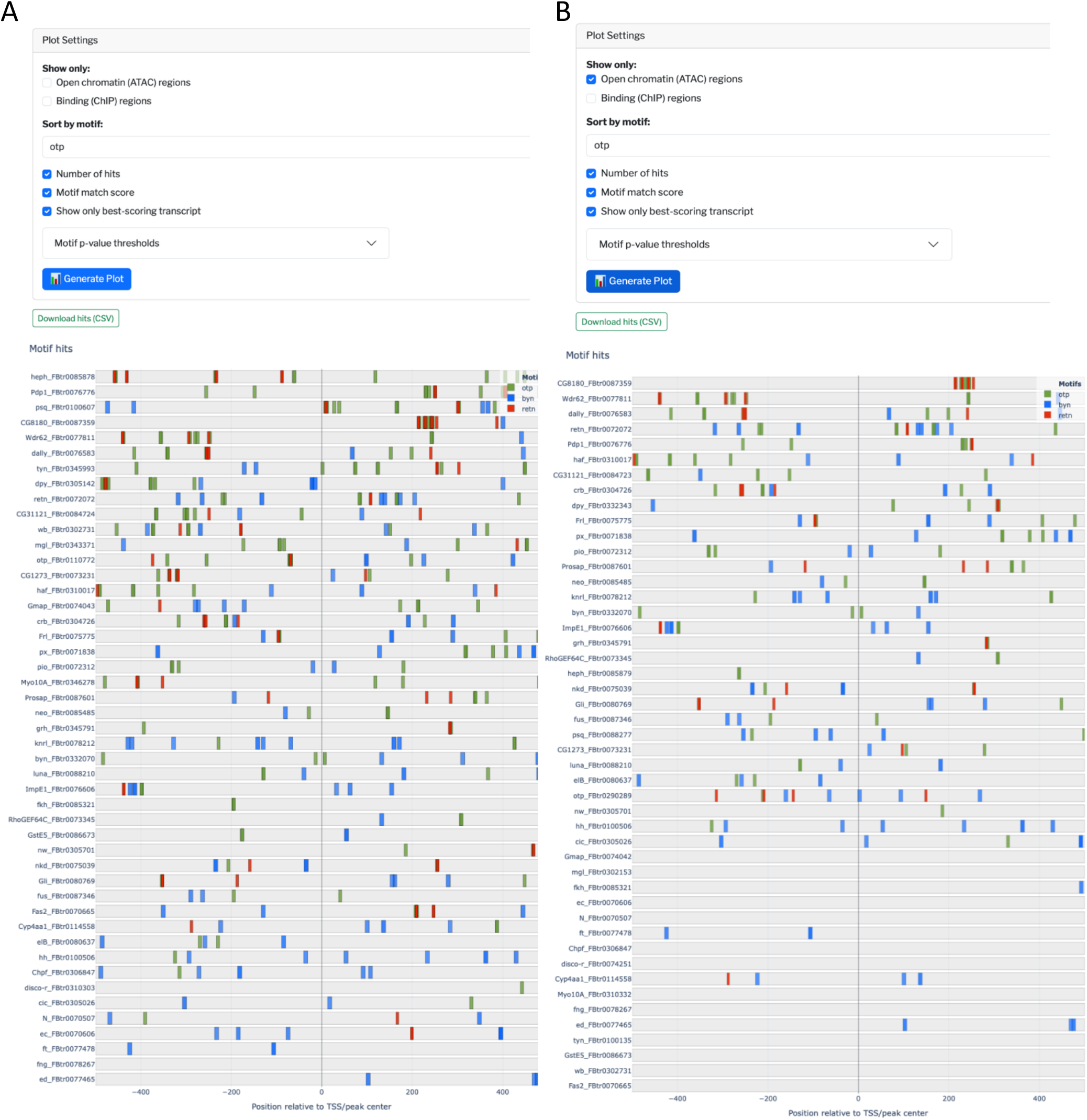
The motif occurrence plot. Motif occurrences of *otp, byn* and *retn* the top 50 DE genes in the hindgut. Plot settings configured to sort the genes by *otp* hits and match score, and showing only the best scoring transcript per gene. The configured filtered hits can be downloaded using the Download hits (CSV) button. Each plot corresponds to the plot settings above it. **(A)** Genes ranked by *otp* occurrences, scored by number of hits and motif match score. **(B)** Plot further filtered for motif hits only in open chromatin regions. Note that motifs not in open chromatin disappear, and the genes are sorted by the remaining motif occurrences.

The last tunable setting is the individual p-value threshold for each motif. Some motifs are more permissive than others and therefore might need further tuning for match stringency, so users have the option to make the p-values for each motif independently tighter. For this, click on the “Motif p-value thresholds” dropdown and drag the slider down the desired p-value. Once all the settings are decided, click on the “Generate Plot” button and the motif plot will be updated accordingly. All the displayed motif hits can be downloaded from the button “Download merged hits”. For the compare mode, users can also individually download hits for both genes list as a zip file from the “Download individual hits” button.

#### Interpreting the motif occurrence plot

The motif occurrence plot contains all the input genes stacked and aligned along the TSS (**Fig 4**). VIDEO accounts for transcription direction and reverses the strand so that all the genes have upstream to the left and downstream to the right of the center line (zero). Each row in the plot corresponds to a gene’s promoter region, and is named in the GeneSymbol_TranscriptID convention, where the transcript ID is unique to a TSS. The plot is interactive, so users are requested to scroll outside the plot region to go further down the list. Double clicking on the zoomed-in plot can revert it to the original layout, and there are also helper buttons to download the image, zoom in, and reset axes at the top right corner of the plot. Hover over any motif hit to see details about the matched sequence, exact coordinates, strand, p-value and match score. A darker motif hit corresponds to a stronger motif match score, and a more transparent hit reflects a weaker match score.

#### Visualizing ATAC-seq and ChIP-seq tracks along with motif hits

The last window of VIDEO is optional and intended for users who wish to visualize ATAC/ChIP-seq signal tracks in the context of the motif hits. To use this tool, users must first upload their BigWig signal tracks of interest (up to three tracks for a plot), give it a name, and pick a color for the track and click Process Tracks. To view the tracks overlaid with the motif hits, select a gene from “Select a peak” dropdown and select the tracks to visualize, and then click on “Generate Overlay”. This will generate a plot with the motif hits on top and the signal tracks aligned below it, along the TSS with the strand accounted for (**Supplemental Fig S1**). The y-axis values are all auto-scaled as a group, and users can visualize where the peaks in the signal are with respect to the motif hits. This plot is also interactive, and hovering over the tracks will display the signal values and chromosomal coordinates.

### 3. Exploring TFs of interest in hindgut tissue

We first sought to use VIDEO explore the transcriptional landscape of the hindgut tissue and identify candidate TFs that make up the regulatory network of hindgut development. Using VIDEO on scRNA-seq data from stage 10-12 hindgut cells, we identified *orthopedia* (*otp*), *brachyenteron* (*byn*), *and retained* (*retn*) (aka *dead ringer*) as the top three DE TFs, followed by other hindgut-expressed TFs including *fork head* (*fkh*), *luna, drum* (*drm*), and *brother of odd with entrails limited* (*bowl*). (**Fig 2C**). We plotted the occurrences of the binding motifs of the top three TFs in the top 50 DE genes (three of which are the TFs themselves) in the hindgut to identify candidate early targets of these TFs.

*otp* encodes a conserved homeodomain TF required for proper hindgut development (Hildebrandt *et al*. 2020), and is strongly expressed in the hindgut as well as in subsets of the CNS (**Supplemental Fig S2**). Its expression in the hindgut is activated by *byn*, where *otp* regulation depends on the number and arrangement of multiple high and low-affinity *byn* motifs (Kusch *et al*. 2002). In the *otp* promoter, VIDEO identified three *byn* motifs downstream of the TSS, along with one *otp* motif (**Fig 4**).

*byn (brachyenteron)* encodes a T-box TF expressed in cells that internalize to form the hindgut, where it acts as a broad regulator of hindgut identity (Singer *et al*. 1996). In the *byn* promoter, VIDEO revealed two *otp* motifs adjacent to the TSS and one *byn* motif downstream. While this arrangement suggests potential autoregulation, and Brachyury orthologs in other species do autoregulate, prior work indicates that *byn* does not autoregulate in *Drosophila* (Singer *et al*. 1996). The observed *byn* motif may therefore represent a conserved but non-functional site, and experimental validation, such as deleting this motif and assaying *byn* expression would help clarify its role.

The third TF recovered in our analysis was *retn*, which encodes an ARID-domain protein that recognizes AT-rich DNA sequences. *Retn* is expressed in the hindgut epithelium and other tissues such as the amnioserosa, ring gland, glia, etc.; within the hindgut epithelium, it is expressed strongly in the boundary cell rows of the large intestine (Iwaki *et al*. 2001; Shandala *et al*. 2002) (**Supplemental Fig S2B**). While its precise role in hindgut development remains unknown, *retn* is required for early embryonic patterning (Shandala *et al*. 1999) and for CNS development (Shandala *et al*. 2003), where it seems to act in combination with other upstream regulators to modulate gene expression.

Not all motif occurrences reflect true binding events, as some may fall outside accessible chromatin regions or arise by chance. To identify motif hits that are more likely to correspond to regulatory activity in the hindgut, we filtered the motif hits using published scATAC data (Wang *et al*. 2025) focusing on hindgut tissue peaks from stage 10-12 embryos. We broadly restricted peaks to those present in at least 50% of hindgut cells (**Supplemental Fig S3**). Under this cutoff, no motifs were detected in the *retn* promoter, suggesting that the filter was too stringent. Because *retn* expression is limited to a subset of hindgut cells, we relaxed the cutoff to include peaks present in at least 25% of the hindgut tissue. This adjustment restored the expected motif hits in the *retn* promoter, confirming that the restricted expression of *retn* correlates with localized chromatin accessibility. Notably, motifs for *byn, otp*, and *retn* were observed in the *retn* promoter, suggesting possible regulatory interactions that merit further testing in mutant or motif-deletion backgrounds. Further reducing the threshold to 10% did not reveal any additional motifs.

Although all the genes in the list are the top differentially expressed in the hindgut based on scRNA annotation (further confirmed by their high expression in the hindgut from in situ hybridization data in BDGP), we found that the promoter regions of 12 genes lacked detectable open chromatin even at the lowest threshold of 10%. This finding seems counterintuitive: highly expressed genes are generally expected to display promoter accessibility, since transcription requires open chromatin. The discrepancy may stem from the technical limitations of scATAC-seq (e.g., the small number of fragments per cell, high sparsity of signal and locus-level dropout), which, although widely used to assay open chromatin, can fail to capture regions where the promoter accessibility is weak, hindered, or transient (Chen *et al*. 2019; Li *et al*. 2021). That said, we were able to identify candidate regulatory events in 75% of the top DE genes in the hindgut, providing a set of hypotheses that can be further tested experimentally.

### 4. Case study: CrebA motif occurrences in SPCGs

To assess whether VIDEO can recover known transcription factor-target gene relationships, we used CrebA as a test case, asking whether its binding motifs are preferentially enriched and stronger in the promoters of secretory pathway component genes (SPCGs) compared to ribosomal protein genes (RpGs). The SPCGs represent a well-established transcriptional target set for the bZIP TF CrebA in the *Drosophila* salivary gland and other secretory tissues (Abrams and Andrew 2005; Fox *et al*. 2010; Johnson *et al*. 2020; Jackson *et al*. 2025). CrebA directly regulates genes encoding secretory machinery, enhancing the secretory capacity of embryonic secretory tissues such as the salivary gland and epidermis. It has also been implicated in dendrite development through upregulation of CopII components (Iyer *et al*. 2013) and in infection tolerance, where its expression in the fat body is induced by the Toll and Imd pathways to activate secretory pathway genes (Troha *et al*. 2018).

For our analysis, we compared 90 SPCGs (Johnson *et al*. 2020) to 82 cytoplasmic ribosomal protein genes (RpGs), which serve as a size-matched gene set that, like the SPCGs, is expected to be found in open chromatin based on the basal levels of these factors being required in all cells. This analysis also addresses the question of whether CrebA also increases translation by boosting expression of RPGs, a regulatory role that has been suggested for the mammalian CrebA orthologue Creb3L2 in the mammalian pituitary gland. The VIDEO workflow for this experiment is described in **Fig 5**. Motif scanning revealed that CrebA motifs were substantially more abundant in the proximal promoters of SPCGs than in RpGs (**Fig 6A**). We tuned the p-value thresholds for both motifs so that the core ACGTG motif was retained in all hits. This adjustment revealed at least one CrebA motif within 500 bp of the TSS in 84% (76/90) of SPCGs, whereas only 61% (51/82) of RpGs had a motif hit in the corresponding promoter proximal regions (**Fig 6B**). Further, among the genes that had at least one motif hit, we examined the density of motif occurrences within 500 bp of the TSS (**Fig 6C**). In SPCGs, nearly three-quarters of motif-positive genes carried multiple hits, with 22% containing three and 33% harboring four or more motifs. By contrast, motif-positive RpGs were skewed toward single occurrences, with almost half (47%) carrying only one motif and a much smaller fraction (18%) with four or more motifs.

**Figure 5.**
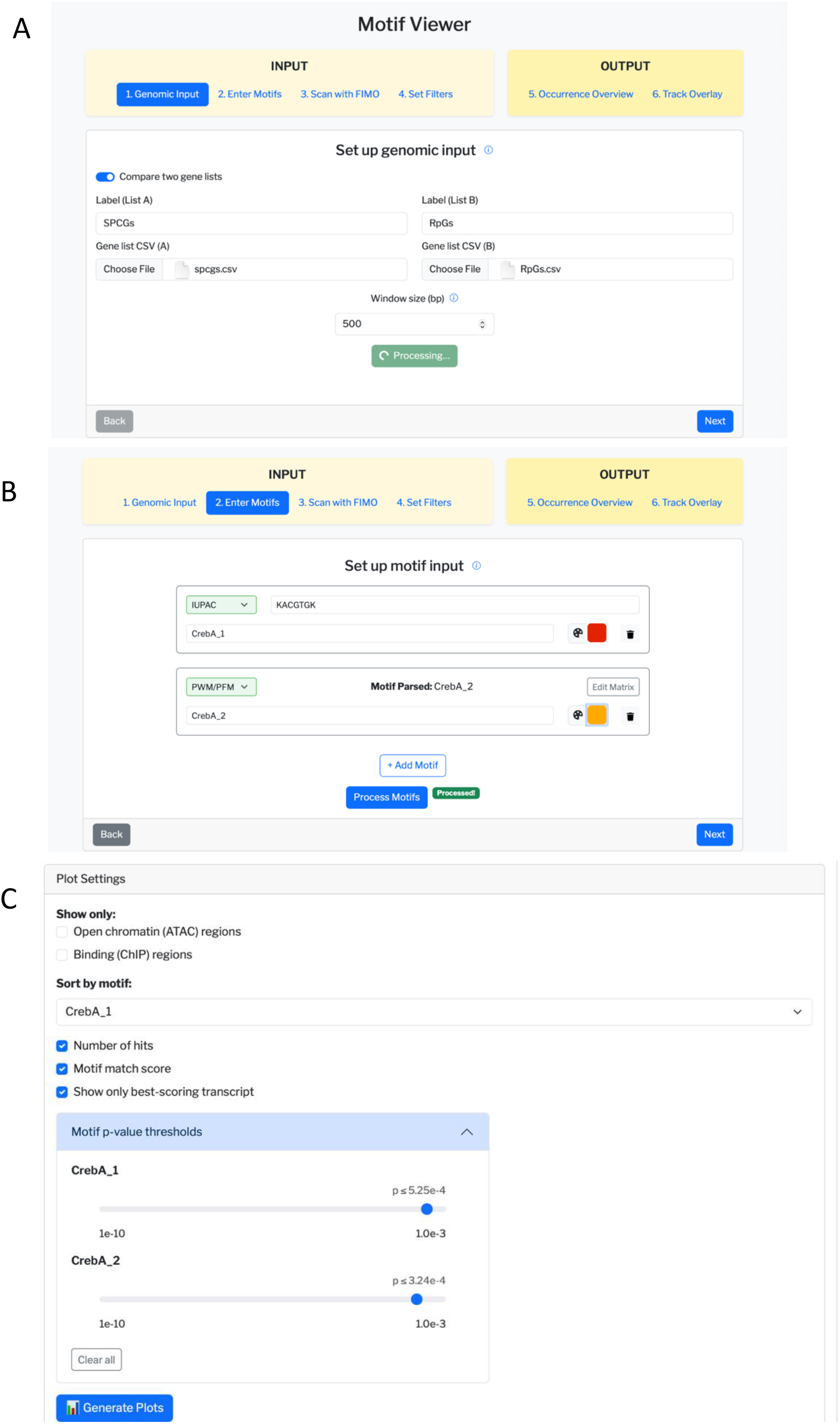
Workflow of CrebA motif occurrence case study. (**A)** In the compare mode, input a gene list of interest (SPCGs) and a control (RpGs) as gene symbols, choose a window size of 500bp, then press process genomic input. **(B)** Input the motifs of interest two slightly different forms of the CrebA motif – the IUPAC consensus sequence from Jackson et al. 2025 (CrebA_1, red) and the PWM from JASPAR 2024 (CrebA_2, yellow) and choose colors for each. **(C)** Adjust plot settings and tune p-value thresholds for individual motifs and then click on Generate Plot.

**Figure 6.**
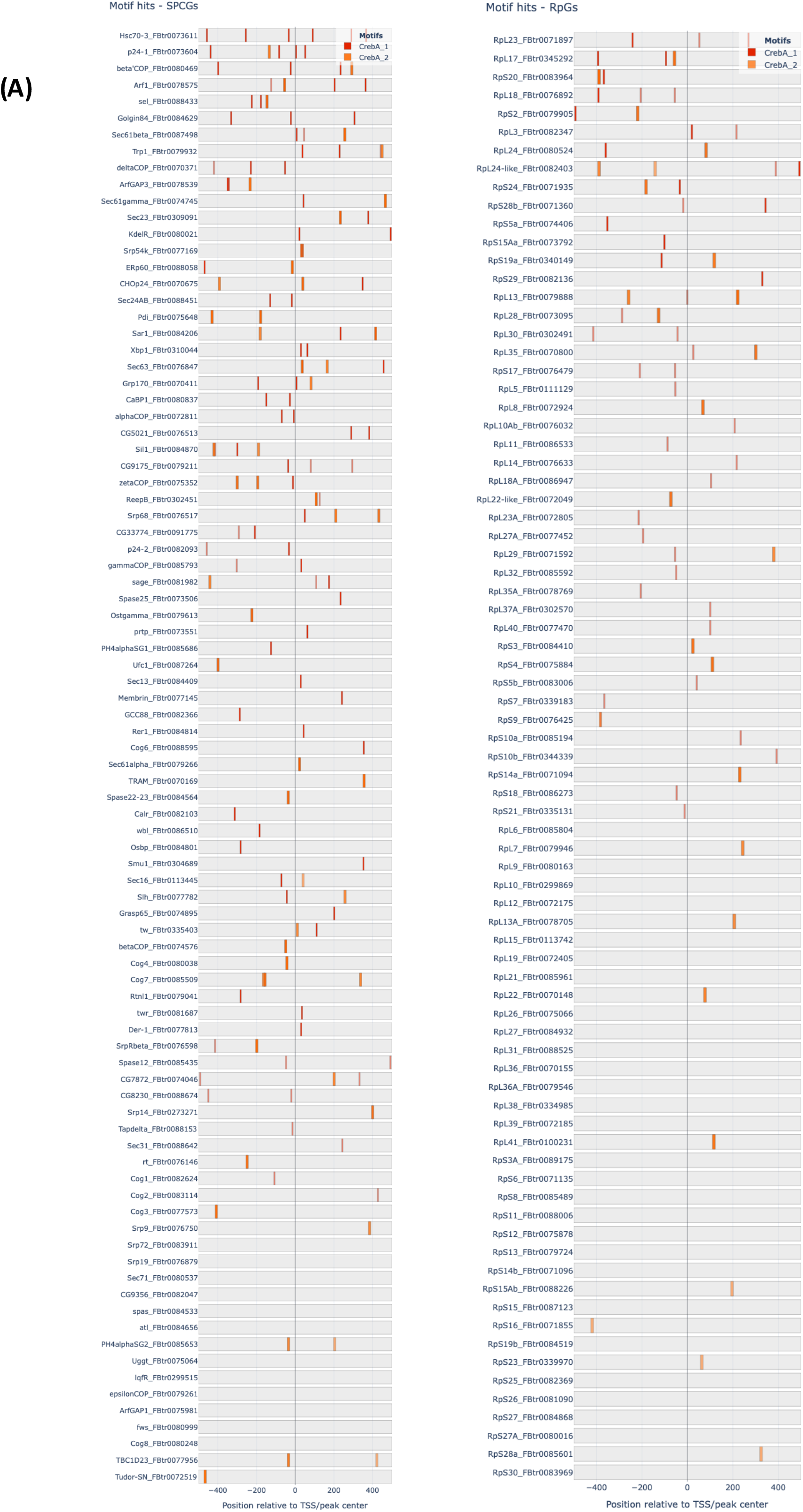
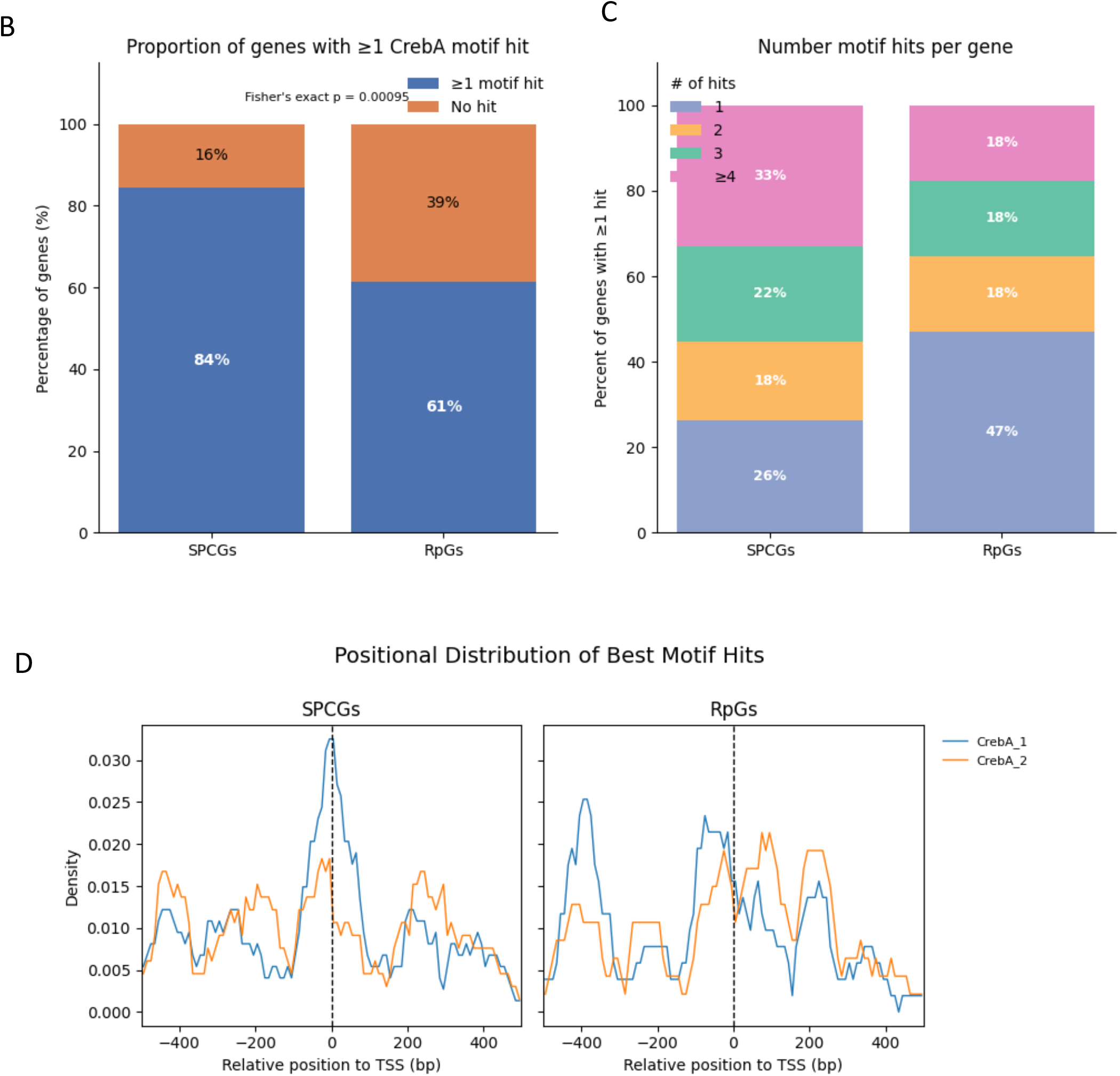
CrebA motifs are more abundant in SPCGs than RpGs. **(A)** Plot shows the occurrence of CrebA motifs in promoter regions of (left) 90 SPCGs and (right) 82 RpGs. Genes are ordered by the number of hits and motif match scores of CrebA_1 motifs. **(B)** The majority of secretory pathway component genes (SPCGs) contain at least one CrebA motif (85%), compared to only 62% of ribosomal protein genes (RpGs). Fisher’s exact test confirms a significant enrichment of CrebA motifs in SPCGs (odds ratio = 3.4, *p* < 0.001). Motif p-value thresholds are CrebA_1: p<5.25e-4 CrebA_2: p<3.24e-4. **(C)** Of the genes that have at least 1 CrebA motif hit, a larger fraction of the SPCGs had more than 4 hits as compared to RpGs. **(D)** The IUPAC sequence version of the CrebA motif (from Jackson et al., 2025, CrebA_1) shows a peak near at the TSS for SPCGs but not for the RpGs, although on quantification, the preference is not significant (KS test p = 0.23). The consensus PWM version of CrebA (from JASPAR 2020, CrebA_2) does not show obvious positional preference for either group.

These results indicate that CrebA binding sites are not only more prevalent in SPCGs than RpGs, but also tend to occur in higher local densities near SPCG promoters. This enrichment of clustered motifs suggests that SPCG regulation may involve stronger or more cooperative CrebA binding compared to RpGs. To test if the CrebA motif occurrences showed any particular positional preference, we plotted the positional distribution of the best hit for each motif in each gene, for both gene lists (**Fig 6D**). The IUPAC consensus sequence (CrebA_1, derived from Jackson et al., 2025) displayed a sharp enrichment immediately upstream of the TSS in SPCGs, suggesting that high-confidence CrebA motifs preferentially cluster at promoter-proximal sites in this gene group, whereas in RpGs they lacked a pronounced positional peak. The more permissive PWM-based motif (CrebA_2, from the JASPAR 2024 database) showed relatively flat distributions in both SPCGs and RpGs, consistent with its broader degeneracy and lower information content. Although these trends qualitatively support the idea that SPCGs may be subject to more direct promoter-proximal regulation by CrebA, quantification using Kolmogorov-Smirnov tests did not detect significant positional biases for either motif (p > 0.05). Thus, while the observed peak near the TSS in SPCGs is suggestive, we cannot rule out the possibility that it arises from stochastic variation rather than actual positional preference.

In addition to motif frequency, motif match quality also differed markedly between the two genes sets. The log-likelihood ratio scores for the motif matches were significantly higher in SPCGs than RpGs, indicating stronger consensus matches among SPCGs (**Fig 7A**). Likewise, log-transformed p-values showed the same trend with SPCGs enriched for significant motif hits compared to RpGs (**Fig 7B**). Interestingly, analysis of motif positions relative to the transcription start sites did not reveal a significant difference between SPCGs and RpGs (**Fig 7C**). This finding is consistent with the positional distribution analysis (Fig. 6D), where CrebA_1 showed a visible promoter-proximal enrichment in SPCGs, but the effect did not reach statistical significance. Together, these data suggest while SPCGs harbor more frequent and higher-quality CrebA motifs, their relative positioning within the 500 bp promoter window is not systematically different from RpGs.

**Fig 7:**
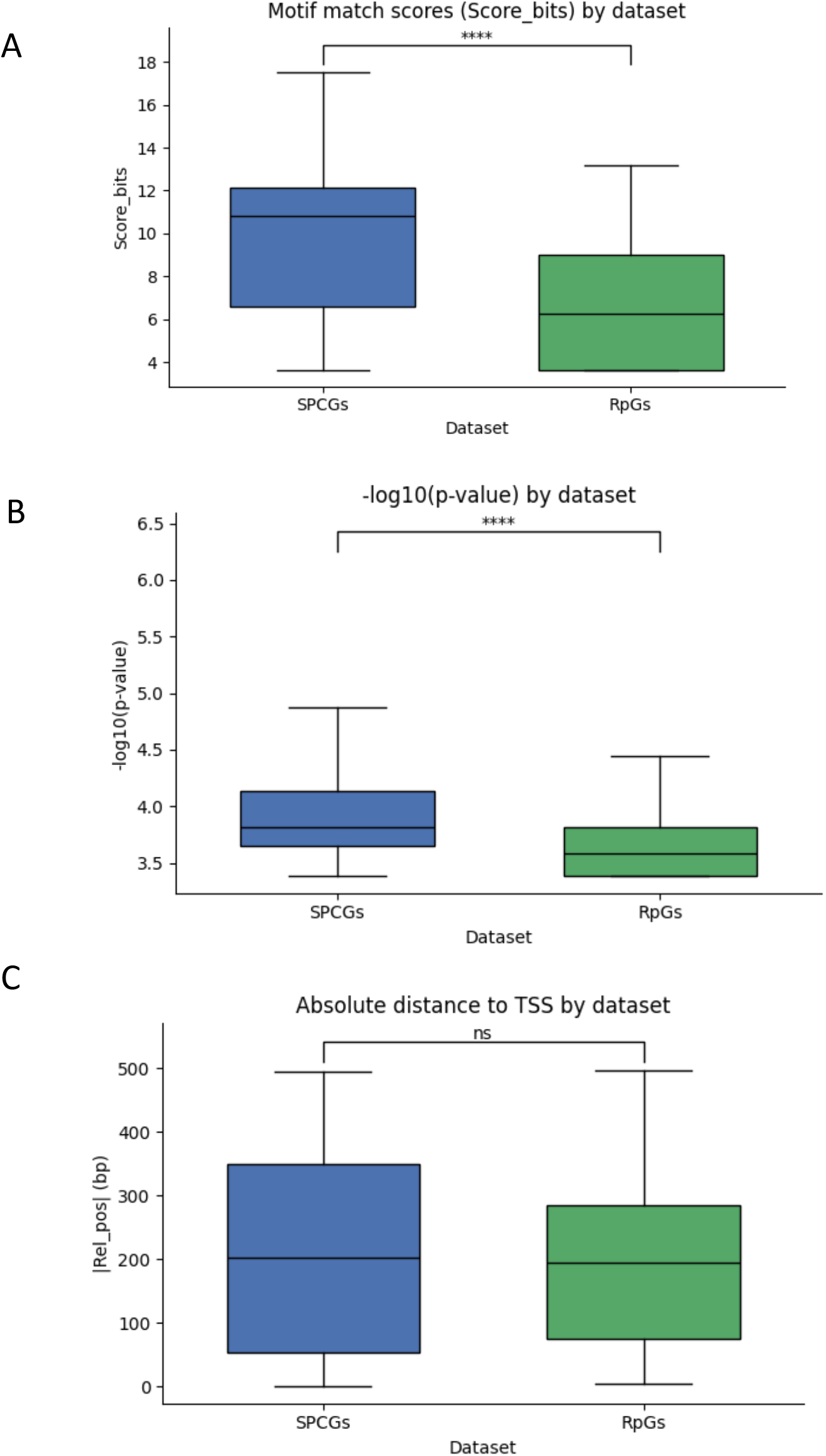
CrebA motif matches are stronger and more significant in SPCGs compared to RpGs. **(A)** Motif match scores (log-likelihood ratio scores) are significantly higher in SPCGs (median = 10.84) than RpGs (median = 6.25; *p* = 1.84e-12). **(B)** Motif match log transformed p-values show the same trend, with SPCGs (median = 3.82) enriched for more significant motif hits (*p* = 3.49e-9) as compared to RpGs (median = 3.58). **(C)** Absolute distance to TSS for all motif hits does not vary significantly between SPCGs (median = 202.0) and RpGs (194.5, p = 0.36). p-values are for all three analyses are from Mann Whitney U-test.

Although the precise positional distribution of motifs remains statistically unresolved, likely due to limited sample size (fewer than 90 genes), the overall enrichment and strength of CrebA motifs in SPCGs align with their established regulatory role. These results confirm that VIDEO accurately captures known biology of CrebA regulation, showing abundance, enrichment and stronger motif matches in SPCGs compared to control. This case study illustrates how VIDEO integrates motif scanning, visualization, and tailored data for statistical comparison into a streamlined user-friendly workflow that validates known transcription factor-gene relationships.

## Discussion

Motif analysis remains a cornerstone for connecting TF binding to regulatory function, but practical barriers continue to limit its accessibility. Existing workflows typically require multiple command-line tools, file conversions and custom scripts, making motif analysis challenging for researchers without strong computational backgrounds. Several visualization tools have emerged, but they often impose restrictions on input formats, motif databases, or integration with genomic datasets.

VIDEO was developed to address these limitations by offering an integrated, web-based platform tailored to *Drosophila* data. It is the first of its kind to facilitate an intuitive side-by-side comparison of user-defined gene lists, incorporates and chooses the best annotated TSSs for each motif based on user-defined metrics such, and allows for ATAC-seq and ChIP-seq data integration (**Table 1**). It also accepts custom motifs and gene list/BED file inputs and presents all the plots in a graphical, interactive format, with downloadable, organized motif hit datasets. These features significantly lower the barrier for motif exploration and visualization compared to existing tools.

**Table 1:** Comparison of VIDEO features to other known motif search and visualization tools.

|  | VIDEO | TFMotifView | MEME-Suite | HOMER |
| --- | --- | --- | --- | --- |
| Online web server | Yes | Yes | Yes | No (command-line) |
| Input format | Gene lists, BED regions, typed genes | BED regions | FASTA sequences | Gene lists, BED regions |
| Compare gene lists | Yes | No | No | No |
| Genome | fly | Human, mouse | Available locally | Available locally |
| Motif database | Custom motifs, JASPAR | JASPAR | Several | Custom |
| View motif occurrences in-app | Yes | Yes | Limited | No |
| ATAC/ChIP data integration | Yes | No | No | No |
| Includes all TSS | Yes | No | No | No |
| Scanning algorithm | FIMO | Motif Information Content | FIMO | HOMER |

The CrebA case study highlights VIDEO’s ability to recapitulate known biology. SPCGs, a well-characterized set of CrebA targets, displayed both higher motif frequencies and stronger consensus matches to CrebA than control RpGs. While the motif positioning relative to the TSS did not differ significantly between the two sets, this could be insightful about CrebA possibly not having a significant preference for a certain position for binding to its target genes provided the binding motif lies within the regulatory window.

Beyond this validation, VIDEO provides a flexible framework that is also suited for hypothesis generation, with helper functions in TF Finder designed to guide users in exploring the transcriptional landscape of the early *Drosophila* embryo. The application of VIDEO to stage 10-12 hindgut data illustrates how visualizing motif occurrences with chromatin accessibility in the right context can be useful for generating hypotheses about gene regulatory networks. The presence of *otp* motifs adjacent to the *byn* TSS, and *byn* motifs downstream of *byn* itself, suggests possible autoregulatory or cross-regulatory interactions. While autoregulation by Brachyury homologs has been demonstrated in other species, it has been excluded for *Drosophila byn* (Singer *et al*. 1996), raising the possibility that these sites are conserved but vestigial or that autoregulation may occur in different developmental contexts. VIDEO thus highlights candidate motifs that can be directly tested by mutation to clarify their functionality. Motif filtering by ATAC-seq data revealed the importance of thresholding stringency when studying genes expressed in rarer cell types, and demonstrates how overly restrictive filters can obscure possibly relevant motifs in genes in specialized subpopulations.

By combining motif scanning, visualization, and data integration into a single user-friendly tool, VIDEO enables broader adoption of motif analysis approaches and expands the potential for researchers to link sequence motifs with regulatory function.

## Supporting information

Supplemental File linked to Supplemental Figures 1, 2, 3

## Acknowledgements

We thank Flybase for the genomic data that was essential for the development and utilization of this tool. We gratefully acknowledge support from the National Institutes of Health (NIH) for support for this research: NIH RO1 DE013899.

