## Supplemental File linked to Supplemental Figures 1, 2, 3 for "VIDEO - Visual Integration of Drosophila Enhancer Organization: A tool for integrating and visualizing chromatin accessibility, in vivo transcription factor binding and motif occurrence in tissue-specific differentially expressed genes"

**Supplemental figure legends**

**Supplemental Figure S1.** *retn* motif track with single cell ATAC peak data. Image shows output of motif occurrence for Otp, Byn, and Rtn consensus binding sites 500 bp upstream and downstream of the *retn* transcription start site (gray bar spanned by colored lines). The graph below shows the open chromatin peaks spanning the same region obtained from single cell ATAC-seq data from stage 10 – 12 hindgut (Wang et al., 2025).

**Supplemental Figure S2. (A)** Expression patterns of top DE TFs in the hindgut, in situ hybridization images from BDGP. **(B)** Drosophila embryos stained with anti-Retn (Dri). Embryos on the left are lateral views, those on the right are dorsal views. Retn is first detected in the region of the hindgut primordia during embryonic stage 10 (top left panel, arrow). From stage 12 – 16, Retn is detected in the nuclei of the boundary cells in the hindgut (all other panels, arrows) and in the distal-most hindgut cells that are found adjacent to the developing Malphigian tubules (bottom two panels, arrowhead).

**Supplemental Figure S3.** Occurrence of *otp, byn,* and *retn* motifs in the top 50 DE genes in the hindgut, filtered by ATAC-seq data (Wang et al., 2025) and sorted by *otp* motif hits. (Left) open chromatin in at least 50% of hindgut cells. (Right) open chromatin in at least 25% of hindgut cells

*retn* is expressed in only a subset of hindgut cells and hence accessible motif hits in the *retn* transcript are recovered when using a lower threshold for ATAC peak calling in the hindgut.
