## Supplemental Table 1 for "VIDEO - Visual Integration of Drosophila Enhancer Organization: A tool for integrating and visualizing chromatin accessibility, in vivo transcription factor binding and motif occurrence in tissue-specific differentially expressed genes"

**Supplemental Table S1.** Abbreviations and definitions for motif representation formats commonly used terms in the paper.

| Abbreviation | Term | Explanation |
|---|---|---|
| PFM | Position Frequency Matrix | A matrix describing a DNA/RNA motif. Each row corresponds to a position in the motif, and each column represents one nucleotide (A, C, G, T). In a PFM, entries are the observed counts of each nucleotide at each position across aligned sequences. |
| PWM | Position Weight Matrix | A matrix describing a DNA/RNA motif. Each row corresponds to a position in the motif, and each column represents one nucleotide (A, C, G, T). In a PWM, these counts are converted to probabilities or log-odds scores, reflecting the likelihood of each base occurring at that position. |
| PCM | Position Count Matrix | A raw count matrix derived from aligned motif instances. Each row is a position in the motif, and each column (A, C, G, T) contains the number of times that nucleotide was observed at that position across the sampled sequences. PWMs and PFMs are typically normalized versions of PCMs. |
| IUPAC | IUPAC consensus sequence | A single consensus sequence that uses IUPAC nucleotide codes to represent base degeneracy. Each letter may correspond to more than one possible nucleotide (e.g., **K = G or T**). While compact, this format captures less information than matrix-based representations because it does not quantify base frequencies. |
| | Consensus Motif | A motif that is a consensus binding site and can be represented in any of the forms above. |
