## Supplementary figures and images for "VIDEO - Visual Integration of Drosophila Enhancer Organization: A tool for integrating and visualizing chromatin accessibility, in vivo transcription factor binding and motif occurrence in tissue-specific differentially expressed genes"

### Supplemental Figure 1

#### Supplemental Figure S1

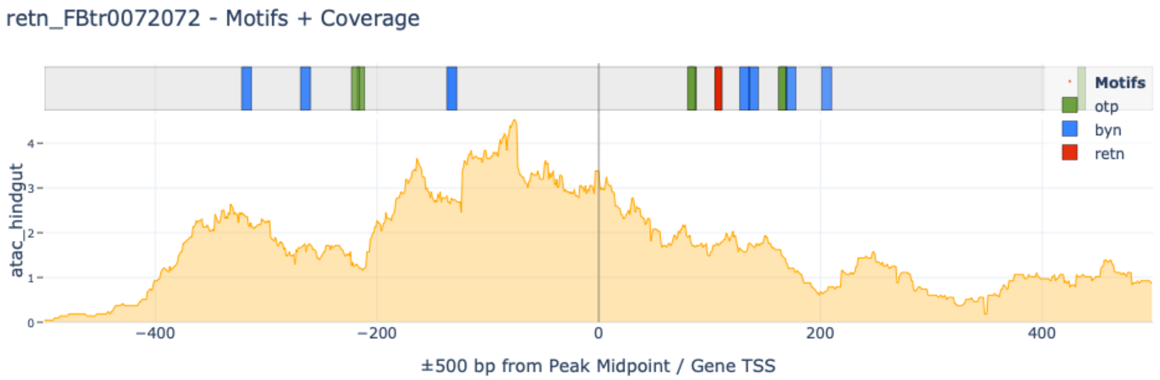

### Supplemental Figure 2

#### Supplemental Figure S3

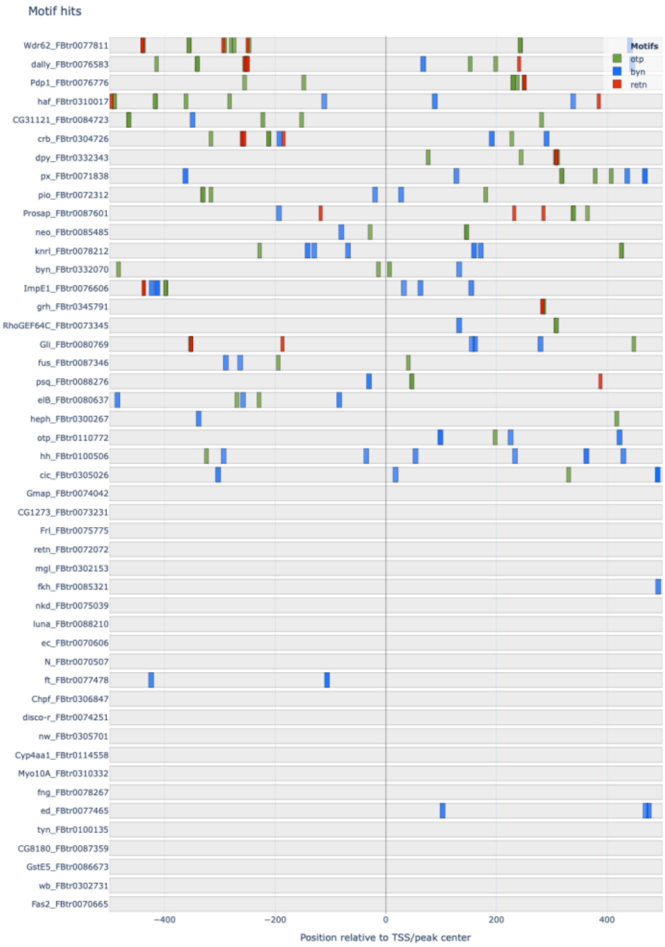


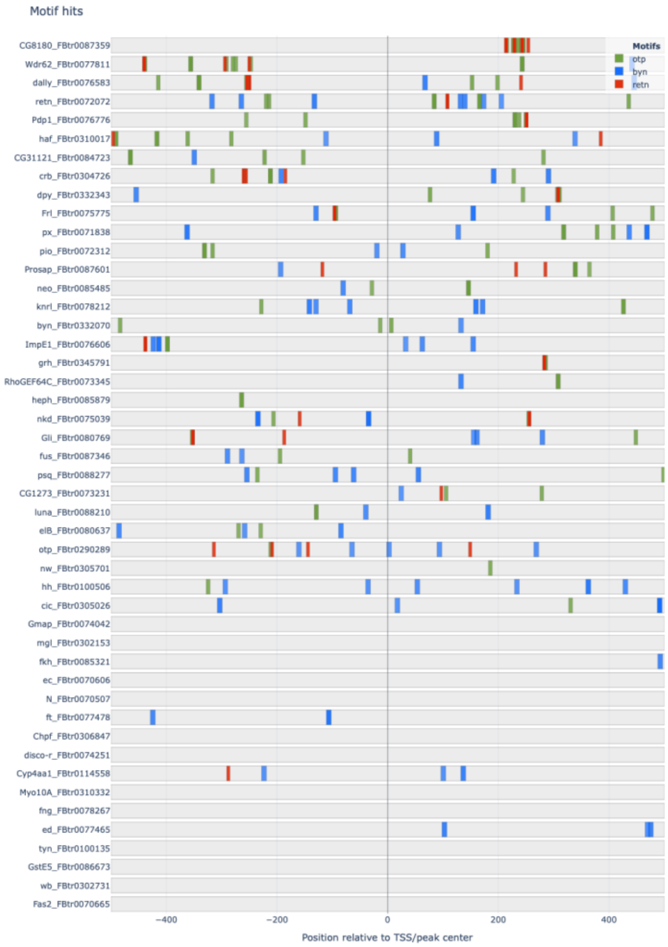
