## Supplemental Figure 3 for "VIDEO - Visual Integration of Drosophila Enhancer Organization: A tool for integrating and visualizing chromatin accessibility, in vivo transcription factor binding and motif occurrence in tissue-specific differentially expressed genes"

### Supplemental Figure S2

(A) Expression patterns of top DE TFs in hindgut

10-12 | 13-16

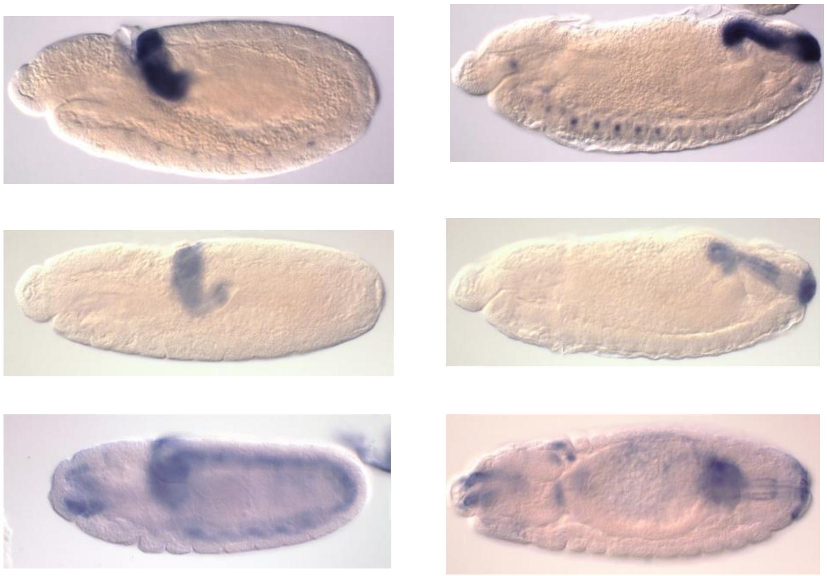

*otp* — strong hindgut, CNS

*byn* — hindgut

*retn/ dri* — hindgut, other tissues

*All in situ images from BDGP*

(B) Retn (Dri) expression in the hindgut

Lateral views | Dorsal views

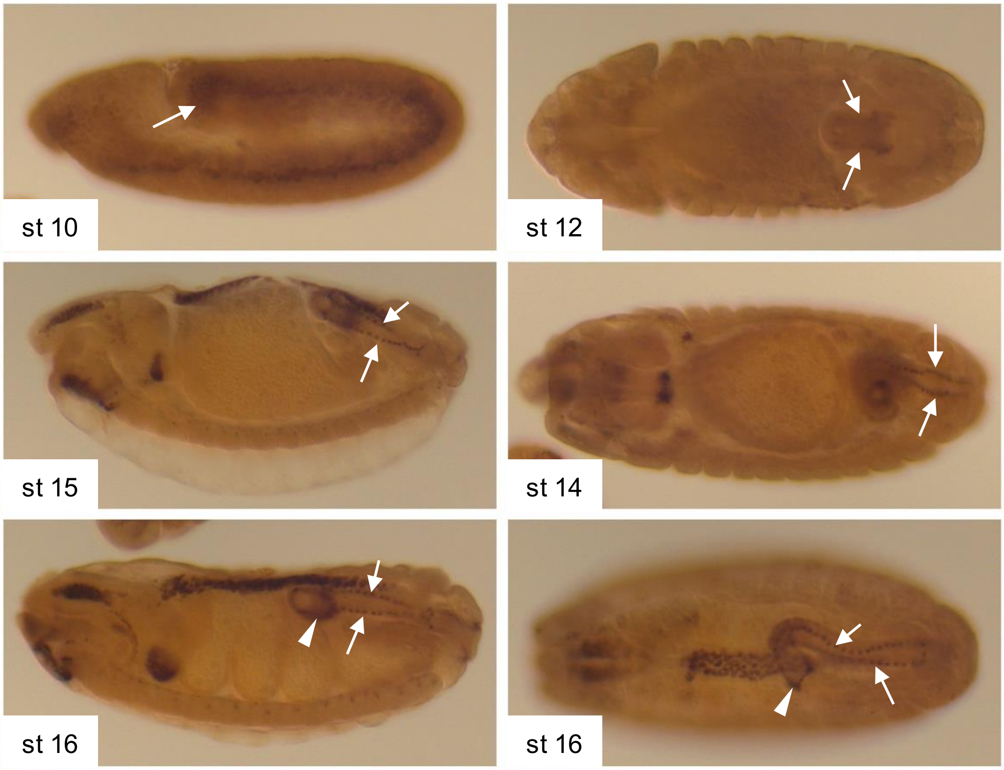
